# An anticipated optimization approach to macromolecular crystallization

**DOI:** 10.1101/620328

**Authors:** Fabrice Gorrec

## Abstract

Crystallization is an essential step for determining macromolecular structures at atomic resolution with X-ray crystallography. Crystals of diffraction quality obtained from purified samples of proteins, RNAs, DNAs and their complexes enable our understanding of biological processes and structure-based design of drugs. Targets of interest for researchers are however increasingly challenging to produce and crystallise. Progress in crystallization methods applicable to limiting amount of sample while increasing the yield of useful crystals are hence urgently needed. In this context, an Anticipated Optimization Approach was investigated. For this approach, it is assumed that samples are highly unstable and will most probably not produce useful crystals (‘hits’). By selecting leads very early, what remains from the initial sample can be used for follow-up optimization experiments. Subsequently, the reproducibility issues caused by sample variability are bypassed. An initial crystallization screen that failed to produce hits becomes a well-suited solubility assay that is a starting point for optimization. The approach was tested with a straightforward and cost-effective protocol developed elsewhere. The yield of useful crystals was increased and accelerated for three targets of pharmacologic studies.

**Synopsis:** This study suggests that the yield of useful crystals obtained from challenging protein samples can be increased by using an initial crystallization screen as solubility assay.

## 1. Introduction

Crystallization of purified proteins, RNAs, DNAs and their complexes is a stochastic process and hence the corresponding screening procedures are mostly empirical (Chruszcz et al., 2008). Essentially, sample variants are screened against hundreds of conditions made of reagents that alter the solubility and stability of macromolecules, the nucleation and growth of crystals, and the main parameters of crystallization experiments (McPherson, 2017). A widespread approach is to apply the technique of vapor-diffusion in nanoliter droplets with several 96-condition initial crystallization screens, which formulation is based on stochastic combination of reagents and former observations (Carter & Carter, 1979; Jancarik & Kim 1991). The underlying problem is the very low yield of diffraction-quality crystals. In addition, optimization experiments (follow-up) are almost systematically required (to produce better quality crystals, or at least enough crystals to thoroughly investigate X-ray diffraction).

The poor solubility and stability of the samples investigated heavily contribute to the difficulties of obtaining useful crystals (Arakawa and Timasheff, 1982; Arakawa and Timasheff, 1985). In this perspective, one can screen a sample against stabilizing buffers/additives before crystallization (Zulauf & D’Arcy, 1992; Jancarik et al., 2004; Ericsson et al., 2006). Nevertheless, there are underlying limitations to pre-crystallization assays, for example the relatively low concentration of sample and reagents required, while crystallization occurs in supersaturated conditions (Asherie, 2004). Besides, most samples purified nowadays will not be stable enough to proceed with a succession of screening assays. With the vapor-diffusion technique, crystals start growing within an hour for some samples, more typically within days, sometimes weeks. Thus, the optimization process often requires frozen aliquots or further sample preparation, which is causing reproducibility issues (Cao et al., 2003). Hypothetically, a way to minimize the issues caused by the challenging nature of the samples would be to select what conditions have to be optimized shortly after initial screening, then use more of the same sample for follow-up experiments.

In this context, an anticipated optimization approach was tested on three protein samples of pharmacologic studies. The first sample was a preparation of an ESCRT complex (‘ESCRT-II’). ESCRT proteins are involved in the narrow membrane necks that are formed during viral, exosomal and intra-endosomal budding from membranes, as well as during cytokinesis and related processes (Schöneberg et al., 2017). The second sample contained a purified Human p53 DNA-binding domain (‘QMp53y220’). The DNA-binding domain of the tumor suppressor p53 is inactivated by mutation in 50% of human cancers (Joerger et al., 2006). The third sample was an HIV-1 capsid hexamer (‘HIV-1-CA’). HIV-1 capsid participates in all infectious steps from reverse transcription to integration site selection (Freed, 2015). Vapor-diffusion nanoliter droplets were set up using a 96-condition sparse matrix screen. Conditions of interest (leads) were selected for follow-up experiments an hour after initial crystallization screening. This, to make use of the samples supposedly still at their peak of stability. Three of the selected initial conditions were randomly chosen to strongly bias follow-up plates according to straightforward and cost-effective protocol developed elsewhere (Birtley and Curry, 2005; St John et al., 2008). When no crystals were readily obtained in the initial screen, other types of leads were selected for follow-up, such as experiments where the sample was visibly soluble; the initial screen was then a pre-crystallization solubility assay (Jancarik et al., 2004). Useful crystals were produced in each case, while the overall process was facilitated and accelerated.

## 2. Material and method

The preparation of the protein samples has been fully described elsewhere: ESCRT-II complex (‘ESCRT-II’), 115 kDa, 7 mg ml^−1^ (Teo *et al.*, 2004); Human p53 DNA-binding domain Y220C mutant (‘QMp53y220c’), residues 94-312, 49 kDa, 6.0 mg/ml (Joerger *et al*., 2004); HIV capsid (‘HIV-1-CA’), 25 kDa, 32 mg ml^−1^ (Price *et al.*, 2014).

The crystallization protocol (for a single follow-up screen) is shown on the Figure 1. The protocol is based on findings made elsewhere (Birtley and Curry, 2005; D’Arcy et al., 2007; St John et al., 2008). The crystallization plates were the 2-lens original MRC plates for sitting droplets (Swissci) pre-filled with 80 μL of conditions in reservoirs (Stock et al., 2005). Droplets were prepared in upperwells of the plates at 20°C using a Mosquito robot (TTP Labtech). In order to minimize the uncontrolled liquid-liquid diffusion in crystallization droplets at the beginning of the experiments (Reiss *et al.*, 2009), the automated dispensing protocol included mixing (settings: 5 ‘mix cycles’, 150 nL ‘mix volume’ and 0.25 mm ‘mix move’). The plates were swiftly sealed with CrystalClear tape (Hampton Research) and then stored at 18°C. Droplets were visualized with a M-80 stereo-microscope (Leica) equipped with a Micropublisher camera (Q-imaging).

**Figure 1.**
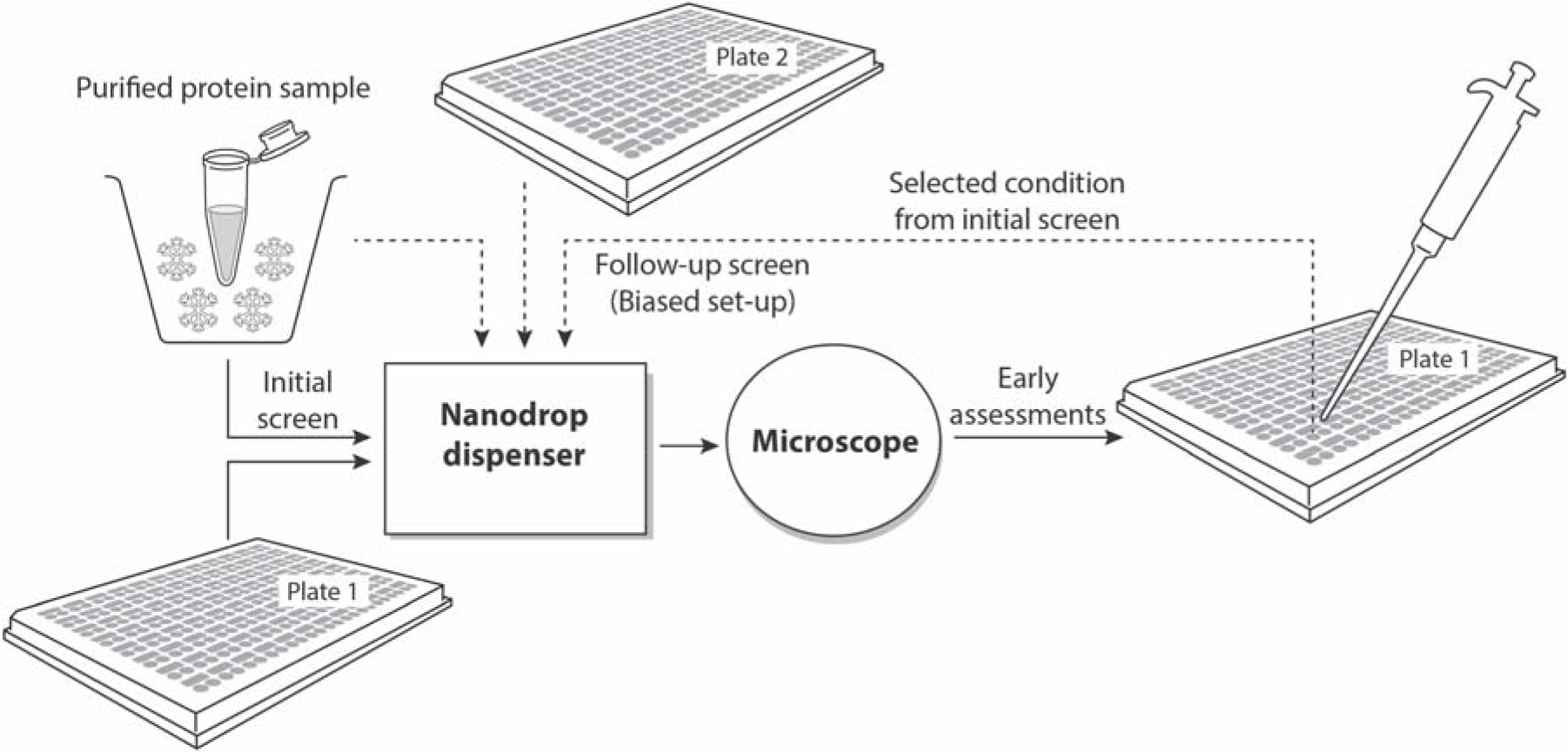
Protocol employed to test the anticipated optimization approach. After early assessments of the droplets in the initial screen (t1 = 1 hr, see Table 1), a selected condition is pipetted out the plate (plate 1). The selected condition enters in the preparation of the follow-up screen droplets (plate 2). The same screen formulation was used in both plates, applying a 4:1 ratio between initial condition and follow-up condition (*i.e*. the follow-up screen is essentially an additive screen). Both plates will be assessed in parallel the following days/weeks (not represented, see Table 2).

For the initial screens (a single plate for each sample), droplets of 500 nL final volume with a 1:1 protein-to-condition ratio were prepared. The leads were selected an hour after setting-up the corresponding droplets. A playing card was associated to each lead and 3 cards were chosen blindfolded to determine which initial conditions will be used to bias the follow-up screens (i.e. 3 additional plates for each sample). Aliquots (75 μL) of the conditions selected for the follow-up screens were pipetted out from the initial plates after a small incision of the seal with a razor blade above the corresponding reservoirs (A piece of tape was used to seal back the corresponding areas of the plates). The initial screen(s) were duplicated (no conditions were extracted from these control plates).

**Table 1.** Types of leads observed an hour after setting up the initial screen (t1) and the three corresponding conditions that were randomly chosen to bias the follow-up screens. For ESCRT-II, three out of the twenty initial conditions that produced clear drops (Figure 2A) were used for follow-up (the sample precipitated in all the other initial conditions). For QMp53y220, two initial conditions producing hypothetical microcrystals (Figure 2C) and one with needle-shaped crystals were picked-up amongst the seven leads. For HIV-1-CA, the three follow-up screens were prepared with initial conditions that produced early microcrystals (dense particles with sharp edges, Figure 2E).

| **Sample** | **Leads** | **Conditions** |
| --- | --- | --- |
| ESCRT-II | 20 clear droplets | A3, A8, B9 |
| QMp53y220 | 5 droplets with hypothetical microcrystals, 2 with needles | C9, D5, D8 |
| HIV-1-CA | 9 droplets with microcrystals, 9 with hypothetical microcrystals, 1 with needles | C7, C8, G6 |

**Table 2.**
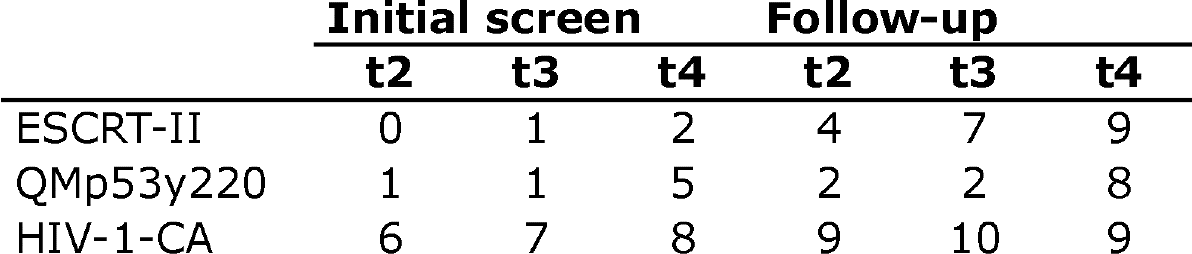
Number of hits observed in the initial screen and the most successful follow-up screen (out of 96 conditions). The different time points are t2:24h, t3:4d and t4:15d. Data from duplicated initial screens were combined (to show the best possible results overall). The most successful initial conditions to bias the follow-up screens were A3 for ESCRT-II, C9 for QMp53y220 and C7 for HIV-1-CA.

Table 1 summarizes the assessments of the initial screens listed in the **Supplementary Material**. The **Supplementary Material** is an Excel file that comprises a suggested nomenclature for scoring droplets, a blank scoring spreadsheet, three scoring spreadsheets with the assessments for the three samples investigated (it was not possible to assess the follow-up plates at t1), the assessments of the control experiments and the formulation of the 96-condition sparse matrix screen used during both initial and follow-up experiments (the ‘LMB crystallization screen’ (Gorrec, 2016), Molecular Dimensions).

For the follow-up screen droplets, the 250 nL of condition were formed with 200 nL of the initial condition and 50 nL of each condition from the follow-up screen. The initial condition to follow-up condition 4:1 ratio was chosen according to work published elsewhere (Birtley and Curry, 2005). A droplet was considered as a hit when it contains well-known single crystals longer than 20 μm and larger than 10 μm.

## 3. Results and discussion

Six out of the nine follow-up screens produced useful crystals. Table 2 shows the corresponding number of hits in the initial screen compare to the most successful follow-up screen (in terms of number of hits) at different time points (starting at t2:24h). The Figure 2 shows light micrographs of droplets for the three samples in the initial screen at t1 and in the most successful follow-up screen at t2.

**Figure 2.**
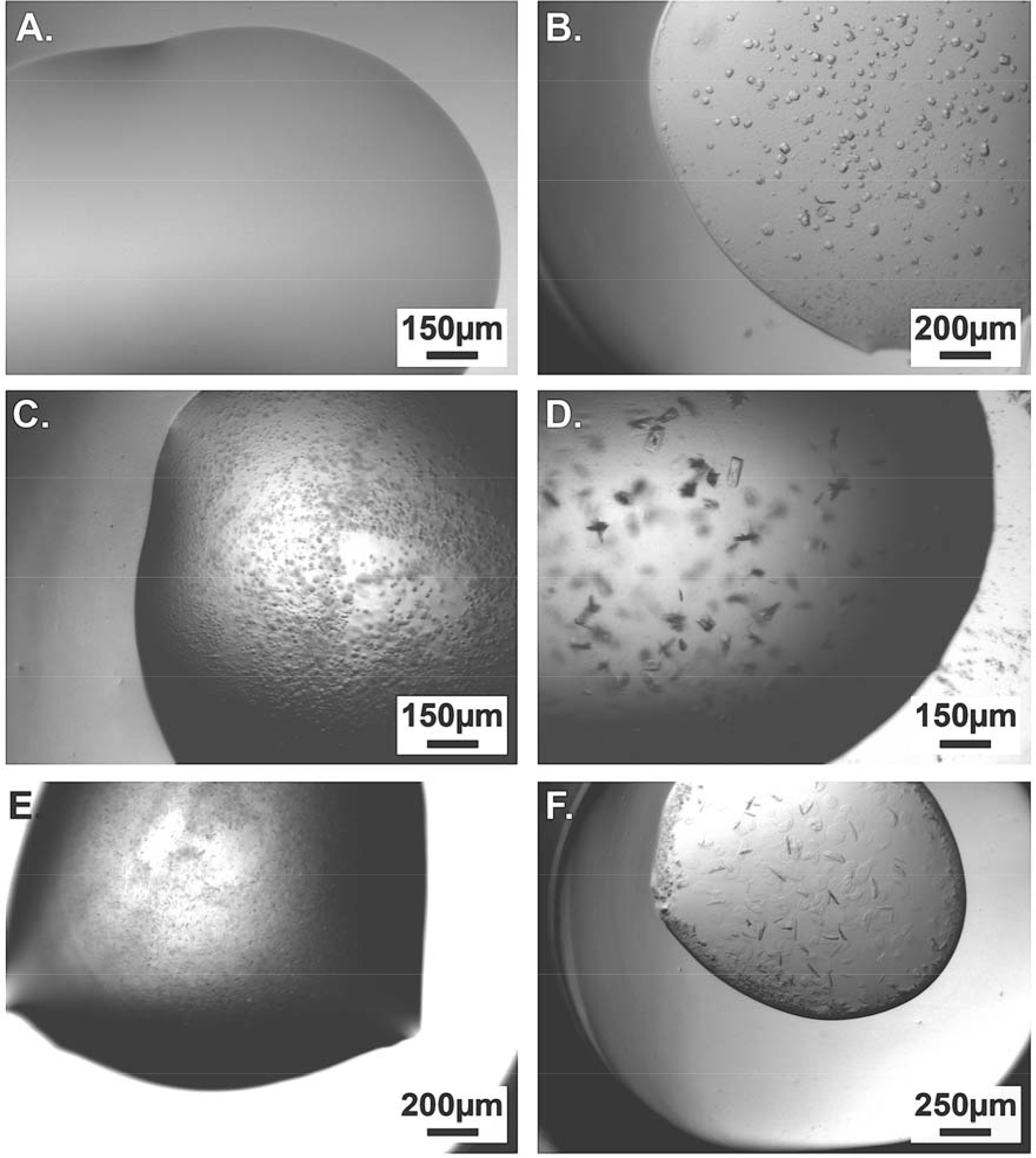
Light micrographs of early leads observed in the initial screens (left) and the corresponding hits observed in follow-up experiments (right). With respectively **A, B**. ESCRT-II against initial condition A3 and later against a mix of condition A3 and D2 as additive (A3:D2). **C, D**. QMp53y22, initial condition C9 and follow-up condition C9:F5. **E, F**. HIV-1-CA, initial condition C7 and follow-up condition C7:F4.

Interestingly, despite the fact that the leads were clear droplets (Figure 2A) for the ESCRT-II sample (and that no hits were observed later in these droplets from the initial screen), two out of three follow-up screens produced hits. The two successful follow-up screens were biased with initial conditions A3 and B9 (**Supplementary Material**). Four hits were observed after 24 hr in the follow-up screen A3 (none in the initial screen).

After 15 days, only 2 hits were observed in the initial screen, 9 hits in the follow-up screen A3. It is worth mentioning that both initial conditions A3 and B9 included an organic volatile reagent as precipitant (propanol and ethanol, respectively) while the condition A8 used to bias the unsuccessful follow-up screen contained a precipitant with radically different properties (ammonium sulfate). The condition A8 had also the most acidic buffer-pH (5.5 compare to 6.2 and 7.1 for A3 and B9 respectively).

In fact, the initial crystallization screen against ESCRT-II was a pre-crystallization solubility assay. New combinations of reagents were generated that induced a more favourable thermodynamics and kinetics to crystal nucleation and growth during the follow-up experiments (Rupp, 2015; Deller et al. 2016). However, the supersaturation required for nucleation to occur might not have been reached during later optimization experiments, because of an insufficient addition of suitable reagents. Hence one could argue that different screen(s) and ratio(s) should be investigated during the follow-up. Ultimately, there are a multitude of other (uncontrolled) parameters to consider. Recent findings show the polymorphic nature of crystal nucleation, growth and structure (Yamazaki et al., 2016; Thompson et al., 2018). Other recent findings show that crystallization experiments can lead to different results with only a slight variation of the protein or the condition according to mechanisms occurring at the very early stage of nucleation (Van Driessche et al., 2018).

For the QMp53y220 sample, the three follow-up screens produced hits in droplets biased with the initial conditions C9, D5 and D8 (which all produced late hits in the initial screen). These three conditions included a high MW polyethylene glycol as precipitant. However, the least successful follow-up screen (D5) had a very basic buffer-pH (9.2 compare to 5.8 and 6.8 for C9 and D8), which probably restricted the yield. At the end, one could argue that, for QMp53y220, the protocol employed here only facilitated an additive optimization screening (Cudney et al., 1994).

For the HIV-1-CA sample, the increase in yield was less pronounced, only one follow-up screen actually worked. The corresponding bias condition (C7) was the only one that contained Magnesium and an acidic buffer-pH (4.6). These two components might be essential to increase the yield of useful crystals of HIV-1-C. In any case, an anticipated optimization seems less relevant when useful crystals are readily obtained in the initial screen. It is then more appropriate to apply a systematic approach to optimization for fine tuning of the initial reagent concentrations and pH (Stock et al., 2005).

## 4. Conclusion

The efficiency of an anticipated optimization approach for macromolecular crystallization was demonstrated. It is particularly well suited for poorly soluble samples that are reluctant to crystallize. The corresponding requirements for the protocol suggested here have been minimized, notably by employing the same screen formulation for solubility assays, initial screen and follow-up experiments. This protocol can hence be easily implemented in most laboratories studying macromolecular structures with X-ray crystallography.

Because the number of experiments to be assessed in a relatively short period of time can be relatively large, and also because some combinations of conditions can produce salt crystals, the anticipated optimization approach would benefit from developments in the field of automated inspection of experiments and assessment of crystals (Wampler *et al*., 2008; Rosa *et al*., 2018).

It would be interesting to apply an anticipated optimization approach to other cases, notably when early gel formation occurs in the initial screen (Van Driessche et al., 2018).

## Acknowledgment

I thank Olga Perisic (MRC-LMB, Cambridge, UK), Andreas Joerger (Institute of Pharmaceutical Chemistry, Frankfurt, Germany) and David Jacques (School of Medical Sciences, Randwick, Australia) for sharing their samples and expertise in assessing the corresponding crystals. Figures were prepared by Jo Westmoreland (LMB Visual Aids). I also thank Michael Wilson (Babraham Institute, UK) for early tips and advices. Although the method presented here can be applied with most of the commercially available plates and screens, I must hereby state a conflicting commercial of interest since the MRC plate and the LMB crystallization screen are commercialized by LifeArc (London, UK) under the MRC intellectual property policy.

## References

Arakawa, T. & Timasheff, S. N. (1982). Biochem. 7, 6545–6542.

Arakawa, T. & Timasheff, S. N. (1985). Biophys. J. 47, 411–414.

Asherie, N. (2004). Methods 34, 266–272.

Birtley, J. R. & Curry, S. (2005). Acta Cryst. D61, 646–650.

Cao, E., Chen, Y., Cui, Z. & Foster, P. R. (2003). Biotechnol. Bioeng. 82, 684–690.

Carter, C. W. & Carter, C. W. (1979). J. Biol. Chem. 254, 12219–12223.

Chruszcz, M., Wlodawer, A. & Minor, W. (2008). Biophys. J. 95, 1–9.

Cudney, R., Patel, S., Weisgraber, K., Newhouse, Y. & McPherson, A. (1994). Acta Cryst. D50, 413–423.

D’Arcy, A., Villard, F. & Marsh, M. (2007). Acta Cryst. D63, 550–554.

Deller, M. C., Kong, L. & Rupp, B. (2016). Acta Cryst. F72, 72–95.

Edwards, A. M., Arrowsmith, C. H., Christendat, D., Dharamsi, A., Friesen, J. D., Greenblatt, J. F. & Vedadi, M. (2000). Nat. Struct. Biol. 7, 970–972.

Ericsson, U. B., Hallberg, B. M., DeTitta, G. T., Dekker, N. & Nordlund, P. (2006). Anal. Biochem. 357, 289–298.

Freed, E. O. (2015). Nat. Rev. Microbiol. 13, 484–496.

Gorrec, F. (2016). Drug Discov. Today 21, 819–825.

Jancarik, J. & Kim, S. H. (1991). J. Appl. Cryst. 24, 409–411.

Jancarik, J., Pufan, R., Hong, C., Kim, S. H. & Kim, R. (2004). Acta D60, 1670–1673.

Joerger, A. C., Allen, M. D. & Fersht, A. R. (2004). J. Biol. Chem. 279, 1291–1296.

Joerger, A. C., Ang, H. C. & Fersht, A. R. (2006). Proc. Natl Acad. Sci. USA, 103, 15056–15061.

McPherson, A. (2008). Protein Sci. 10, 418–422.

McPherson, A. (2017). Protein Crystallography. Protein Crystallization. Humana Press, New York NY. Methods Mol. Biol. 1607, 17–50.

Newman, J., Xu, J. & Willis, M. C. (2007). Acta Cryst. F63, 826–832.

Price, A. J., Jacques, D. A. McEwan, W. A., Fletcher, A. J., Essig, S., Chin, J. W., Halambage, U. D., Aiken, C. & James, L. C. (2014). PLoS Pathogens, 10,e1004459. https://doi.org/10.1371/journal.ppat.1004459

Reis, N. M., Chirgadze, D. Y., Blundell, T. L. & Mackley, M. R. (2009). Acta Cryst. D65, 1127–1139.

Rosa, N., Ristic, M., Marshall, B. & Newman, J. (2018). Acta Cryst. F74, 410–418.

Rupp, B. (2015). Acta Cryst. F71, 247–260.

Schöneberg, J., Lee, I-H., Iwasa, J. H. & Hurley, J. H. (2017). Nat. Rev. Mol. Cell Biol. 18, 5–17.

St John, F. J., Feng, B. F. & Pozharski, E. (2008). Acta Cryst. D64, 1222–1227.

Stock, D., Perisic, O. & Löwe, J. (2005). Prog. Biophys. Mol. Biol. 88, 311–327.

Teo, H., Perisic, O., González, B. & Williams, R. L. (2004). Dev. Cell, 7, 559–569.

Thompson, M. C., Dasciob, D. & Yeates, T. O. (2018). Acta Cryst. D74, 411–421.

Trakhanov, S. & Quiocho, F. A. (1995). Protein Sci. 4, 1914–1919.

Van Driessche, A. E. S., Van Gerven, N., Bomans, P. H. H., Joosten, R. R. M., Friedrich, H., Gil-Carton, D., Sommerdijk, N. A. J. M. & Sleutel, M. (2018). Nature, 114, 89–94.

Wampler, R. D., Kissick, D. J., Dehen, C. J., Gualtieri, E. J., Grey, J. L., Wang, H-F., Thompson, D. H., Cheng, J-X. & Simpson, G. J. (2008). J. Am. Chem. Soc. 130, 14076–14077.

Yamazaki, T., Kimura, Y., Vekilov, P. G., Furukawa, E., Shirai, M., Matsumoto, H., Van Driessche, A. E. S. & Tsukamoto, K. (2016). Proc. Natl Acad. Sci. USA, 114, 2154–2159.

Zulauf, M. & D’Arcy, A. (1992). J. Cryst. Growth, 122, 102–106.

